# The confidence code: parietal alpha oscillations turn expectations into beliefs

**DOI:** 10.64898/2026.05.09.723995

**Authors:** Luca Tarasi, Anna Pasini, Domenico Romanazzi, Margherita Covelli, Vincenzo Romei

## Abstract

Metacognition, the ability to evaluate whether one’s decisions are correct, is crucial for adaptive behavior. However, confidence judgments are not bias-free: they can be systematically modulated by predictive cues and beliefs, favoring expectation-consistent judgments, while leaving metacognitive precision unchanged. Here, we identify parietal alpha activity as the causal mechanism linking expectations to metacognitive bias. In Study 1, EEG was recorded while 75 participants performed a visual detection task with symbolic cues signaling target probability. Cues induced a metacognitive bias in confidence without altering metacognitive sensitivity, and cue-driven alpha modulation over the right parietal cortex predicted the magnitude of this bias. In Study 2 (N = 88), continuous theta-burst stimulation (cTBS) over the right parietal cortex abolished cue-induced alpha modulation, thereby selectively reducing metacognitive bias, while sham stimulation had no effect. Together, these findings demonstrate that parietal alpha-mediated gain control causally shapes metacognitive judgments, revealing an oscillatory code for predictive metacognition.

## Introduction

Human perception is often described as an integrative mechanism, where expectations merge with sensory evidence to build a coherent experience of reality^1^. In this framework, perceptual decisions emerge from such interaction as the brain’s optimal inference based on the environment’s available information. Critically, the perceptual process does not end with this initial decision but continues with a metacognitive evaluation assessing its precision as well as its coherence with the expected outcome^2,3^. This higher-order computation gives rise to confidence judgments, allowing individuals to adaptively regulate subsequent behavior based on their perceived level of certainty or uncertainty^4,5^.

Within the Signal Detection Theory framework (SDT)^6^, perceptual and metacognitive judgments share a common logic: both involve distinguishing signal from noise. At the perceptual level, sensitivity (*d’*) represents the individuals’ ability to discriminate between relevant sensory signals and background noise, while the criterion (*c*) reflects their tendency (i.e., bias) toward a particular decision. Similarly, metacognitive sensitivity (*Meta-d’*) reflects the individuals’ ability to recognize whether their perceptual decisions were correct, whereas the metacognitive criterion (*Meta-c*) captures a general bias toward reporting higher or lower confidence i.e., a policy mapping internal decision variable onto confidence, independently of metacognitive sensitivity.

Despite extensive research on perceptual decision-making, the mechanisms by which prior expectations shape confidence judgments remain unknown. Indeed, while it is well-established that expectations bias perceptual decisions by shifting the decision criterion^7,8^, their impact on metacognition has been comparatively underexplored^9–11^.

Under the assumption of bounded cognitive resources, confidence judgments would not always result from a perfect metacognitive evaluation of perceptual correctness. To avoid unnecessary cognitive effort and to maintain a form of homeostatic balance, our internal representation could be strongly shaped by information that aligns vs. not aligns with our beliefs and expectations. This adaptive mechanism - commonly referred to as confirmation bias^12^ - allows individuals to preserve a sense of perceptual self-consistency by biasing the metacognitive criterion in an expectation-like fashion.

Notably, the neural mechanisms underlying these processes are not explored yet. Recent electrophysiological studies have demonstrated that alpha oscillations are consistently associated with metacognitive evaluations, as indicated by negative correlations between spontaneous pre-stimulus alpha amplitude and subjective confidence judgments^13,14^. Crucially, expectation-induced biases in perceptual decisions correlate with modulations of pre-stimulus alpha amplitude in parieto-occipital regions^8,15^. Nevertheless, it remains unclear whether prior-induced alpha modulation plays a causal role in the integration of probabilistic prior information into confidence judgments.

To address these questions systematically, we conducted a first experiment using a signal-detection framework to assess the effect played by expectation-like information on metacognitive sensitivity and bias. Participants completed a detection task in which a probabilistic cue indicated the likelihood of the upcoming target. Participants were asked to report the presence or absence of the target (Type-I response) and to rate their confidence in this decision (Type-II response) on a 4-point Likert scale. We expect prior information to selectively modulate the metacognitive criterion rather than metacognitive sensitivity. Additionally, electroencephalography (EEG) activity was recorded during task performance to characterize the oscillatory signatures underlying prior-induced confidence biases. We hypothesize that prior-induced modulation of alpha rhythms is directly associated with shifts in the metacognitive criterion, predicting that the magnitude of cue-dependent alpha modulation will be associated with a larger metacognitive bias.

At the anatomical level, we propose that the parietal cortex (PC) is a key neural hub for integrating prior expectations into metacognitive judgments. Previous studies have indicated that PC activity is involved in the integration of prior knowledge^16–18^, as well as sensorimotor integration^19^, decision-making under uncertainty^20^ and metacognitive processes^21^. Crucially, seminal studies have linked PC engagement to the regulation of alpha-band rhythms ^22–24^. For instance, Capotosto and colleagues^22^ demonstrated that inhibitory stimulation of PC shifted alpha-band dynamics, consistent with a role of PC in shaping cortical excitability.

Building upon these findings and informed by the results from Study 1, we conducted a second experiment, combining EEG and continuous Theta Burst Stimulation (cTBS) to causally test PC’s causal involvement in integrating prior expectations into metacognitive judgments. Specifically, we hypothesize that inhibitory PC stimulation will temporarily disrupt the participants’ ability to modulate alpha oscillations according to probabilistic cues. Consequently, this should reduce the cue-dependent integration of prior information into confidence judgments, thereby diminishing the associated metacognitive bias. Importantly, this effect is expected to be specific to metacognitive bias, with no change in metacognitive sensitivity

## RESULTS

### Study 1

Seventy-five participants completed a basic visual detection task (Figure 1A). On each trial, a checkerboard appeared in the lower-left visual field. The checkerboards either included isoluminant gray circles within their cells (target trials) or were empty (catch trials). Participants were instructed to use the keyboard to indicate whether the target was present. A symbolic cue preceded each checkerboard, indicating the probability of target presence. Three types of cues were used: a high-probability cue (67%), a low-probability cue (33%), and a neutral-probability cue (50%) indicating equal probability of presence and absence. The actual presence of the targets matched the probabilities signaled by the cues, and participants were explicitly informed about that.

**Figure 1.**
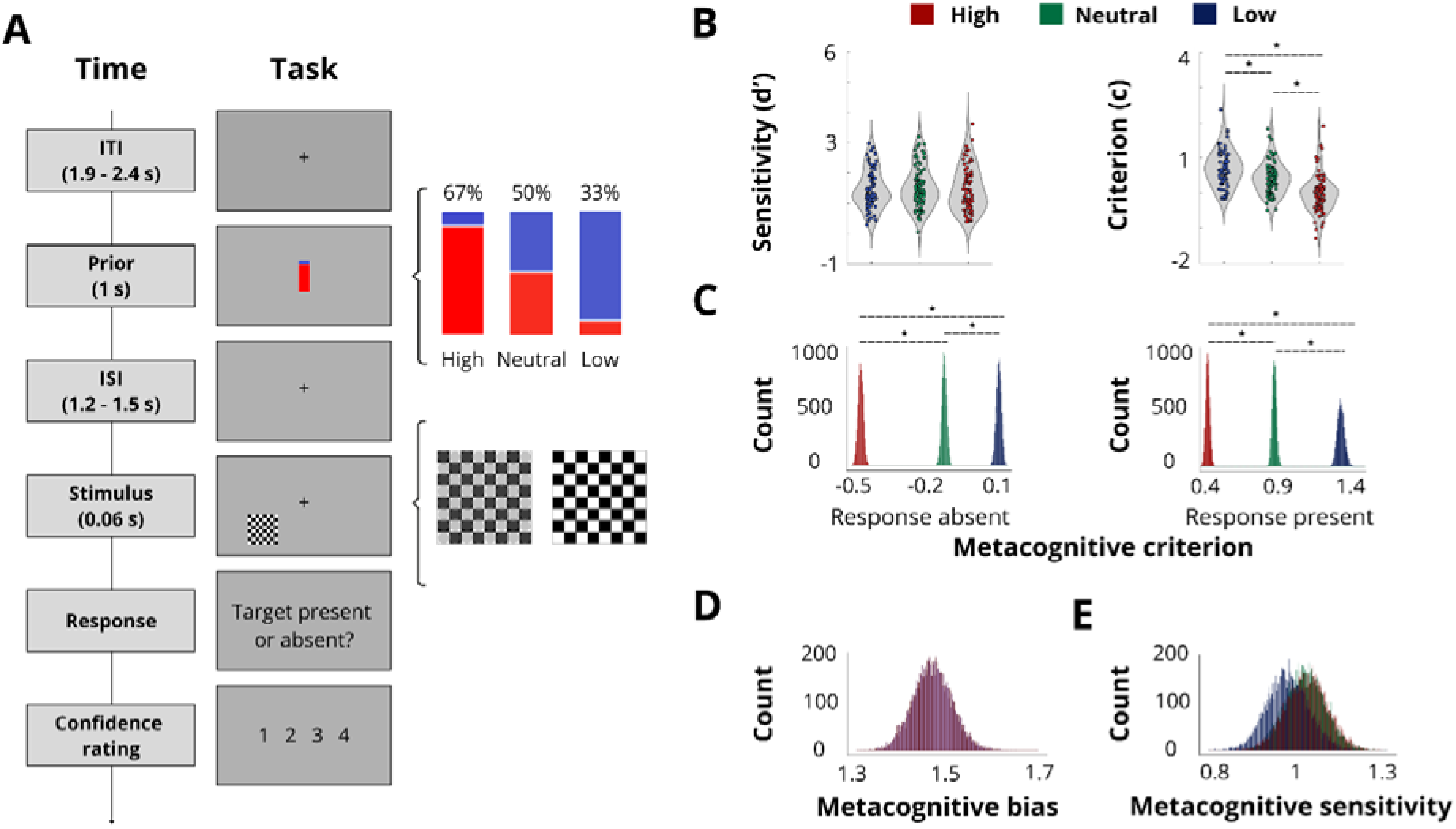
Experimental task and Study 1 results. *A.* Experimental design. EEG data were collected during a simple visual detection task. Each trial began with a fixation cross, followed by the appearance of a probabilistic cue at the center of the screen. Subsequently, a checkerboard containing (or not) grey circles at the titrated contrast within it appeared in the bottom-left of the screen for 60 ms. At the end of each trial participants were asked to indicate how confident they were about their response. Confidence was rated using the keyboard keys “1,” “2,” “3,” and “4” corresponding respectively to four levels of confidence: not confident at all (1), slightly confident (2), moderately confident (3), and highly confident (4). The cue was a rectangle with its bottom colored red and its top colored in blue. The percentage of the red shading to the entire rectangle indicated the target probability. Three cue levels were used: the high and low probability cues indicated the probability of the target presence of 67% and 33% respectively, while the neutral cue predicted the target presence and absence equally (50%). *B.* Type I Signal detection theory. Sensitivity (d’) and Criterion (c) indices are represented separately for trials preceded by high- (in red) low- (in blue), or neutral-probability (in green) cues. The Y-axis represents the values of the two parameters, and each circle corresponds to a subject. Prior information did not affect sensitivity. In contrast, the probabilistic cue significantly influenced the decision criterion: a more liberal criterion was adopted in trials preceded by high probability cue, compared to both neutral and low probability cue. Conversely, a more conservative criterion was adopted in trials preceded by the low probability cue relative to the neutral cue. *C.* Metacognitive criterion (Meta-c). Meta-c index is represented separately for trials preceded by high-, low-, or neutral-probability cues for type-I responses (absent, present). The Y-axis represents the count of observations across the HMeta-d’ distribution, the X-axis represents the most probable HMeta-d’ values. A Meta-c value close to 0 indicates a more liberal criterion (i.e., tendency to give high-confidence ratings), whereas a Meta-c value farther from 0 indicates a more conservative criterion (i.e., tendency to give low-confidence ratings). When a type-I “present” response was given in trials preceded by a high-probability cue, participants adopted a more liberal Meta-c relative to trials preceded by neutral-probability cue and low-probability cue in which participants adopted a more conservative criterion relative to the neutral condition Conversely, when a type-I “absent” response was given in trials preceded by a high-probability cue, participants adopted a more conservative Meta-c compared to trials preceded by a neutral-probability cue, and low-probability cue in which participants adopted a more liberal criterion relative to the neutral condition. *D.* Metacognitive bias index. Higher values indicate greater reliance on probabilistic information when making confidence judgments. *E.* Metacognitive sensitivity (Meta-d’). Meta-d’ index is represented separately for trials preceded by high-, low, or neutral-probability cues. Higher values correspond to greater metacognitive ability. Prior information did not affect metacognitive sensitivity.

### Prior information shapes decision-making strategies

To investigate the effect of expectancy cue on perceptual decision-making, the type I SDT indices *d*’ and *c* were calculated separately (Figure 1B). The conducted analysis showed that cue affected criterion (F_2,148_ = 84.49; p < 0.01; η_p_^2^ = 0.53) but not sensitivity (F_2,148_= 1.38; p > 0.25; η_p_^2^ = 0.02). Specifically, the participants adopted a more liberal criterion in trials preceded by high-probability cue (mean ± SEM = −0.02 ± 0.06) relative to trials preceded by neutral-probability cue (0.42 ± 0.05; t_74_ = −9.15; p < 0.01) and low-probability cue (0.69 ± 0.06; t_74_ = −9.72; p < 0.01), in which participants adopted a more conservative criterion relative to the neutral condition (t_74_ = 7.02; p < 0.01). Overall, the behavioral results indicate that the experimental paradigm used was able to manipulate response bias without affecting other decision-making parameters.

### Prior information shapes confidence criteria but not metacognitive sensitivity

To investigate the effect of the expectancy cue on confidence judgments, the type II SDT indices *Meta-d*_ and *Meta-c* were calculated separately for trials preceded by high-, low-, and neutral-probability cues using *HMeta-d’* toolbox^25^. The analysis showed that prior probabilistic information shapes *Meta-c* (Figure 1C). Specifically, when a type-I “present” response was given in trials preceded by a high-probability cue, participants adopted a more liberal *Meta-c* (mean = 0.45) relative to trials preceded by neutral-probability cue (mean = 0.85; HDI_high-mid_ [–0.43, –0.36]) and low-probability cue (mean = 1.25; HDI_high-low_ [–0.86, –0.76]) in which participants adopted a more conservative criterion relative to the neutral condition (HDI_low-neutral_ [0.35, 0.45]).

Conversely, when a type-I “absent” response was given in trials preceded by a high-probability cue, participants adopted a more conservative *Meta-c* (mean = 0.14) compared to trials preceded by a neutral-probability cue (mean = –0.26.; HDI_high-neutral_ [–0.44, –0.37]), and low-probability cue (mean = –0.53; HDI_high-low_ [–0.71, –0.63]) in which participants adopted a more liberal criterion relative to the neutral condition (HDI_low-neutral_ [0.23, 0.30]). Furthermore, the analysis revealed that the probabilistic cue does not affect *Meta-d’* (Figure 1E; mean low-probability = 1.05; neutral-probability = 1.05; high-probability = 0.99; HDI_high-low_ [–0.22, 0.11]; HDI_high-neutral_ [–0.21, 0.12]; HDI_neutral-low_ [–0.17, 0.17]). These findings suggest that the prior information conveyed by the cue does not modulate the participant’s metacognitive sensitivity but rather induces a metacognitive bias in evaluating the correctness of their choice. Specifically, this effect was congruence-dependent: the same cue modulated metacognitive bias differently depending on whether the participant’s perceptual response was congruent or incongruent with it. Finally, a global Metacognitive bias index was calculated (see Methods) (Figure 1D), with an average value of 1.47 (HDI [1.41, 1.53]). Increasing values in this metric reflect an enhanced weighting of probabilistic information in the formation of confidence judgments.

### Pre-stimulus alpha dynamics encode expectancy information

To examine whether pre-stimulus electroencephalographic activity over posterior electrodes was influenced by the probabilistic information provided by the cue, a non-parametric cluster-based permutation test across the time × frequency domain was conducted (Figure 2A). Specifically, the spectral amplitude of trials preceded by a high-probability cue against those preceded by a low-probability cue was contrasted, focusing on the 500-millisecond interval before stimulus onset and covering a broad frequency range from 4 to 40 Hz. This approach allowed the identifying spatiotemporally contiguous regions of significant difference without assuming parametric distributions. The analysis revealed a reliable modulation in the alpha band, such that high-probability cues were associated with a stronger suppression of pre-stimulus posterior alpha amplitude compared to low-probability cues (p < 0.01). This finding indicates that probabilistic expectations selectively influenced anticipatory neural dynamics prior to sensory processing. Subsequently, a topographical analysis was performed to examine the spatial distribution of the observed pre-stimulus effects (Figure 2A). This involved contrasting high-versus low-probability cue trials for each electrode in the alpha frequency range (∼7-14 Hz) within a 400-millisecond pre-stimulus interval. The topographic cluster-based permutation test revealed a spatially specific modulation of alpha activity over right parieto-occipital electrodes.

**Figure 2.**
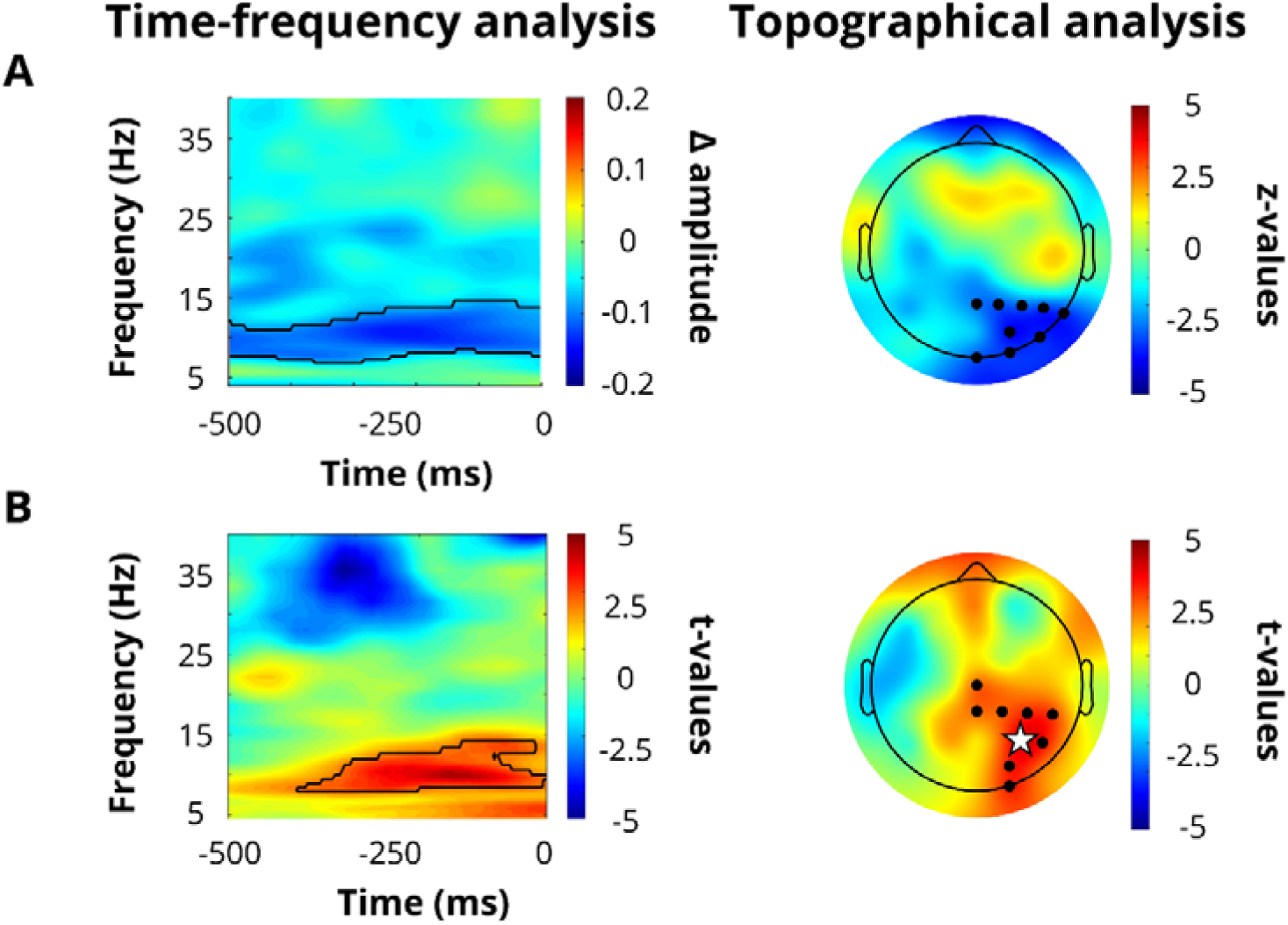
Parietal alpha dynamics predict metacognitive bias. A (left). Time–frequency map of the pre-stimulus (−500 to 0 ms; X-axis) amplitude difference between trials preceded by high- and low-probability cue (Δ amplitude), recorded in posterior electrodes. Time 0 corresponds to stimulus onset. Black contours indicate statistically significant clusters. The Y-axis represents the frequency (4 - 40Hz). There is a suppression in pre-stimulus alpha amplitude in high-probability trials compared to low-probability trials. A (right). Topography of the differential activations in the alpha-band (∼7-14 Hz) between trials preceded by high- and low-probability cue during the pre-stimulus window (−400 to 0 ms). Black dots indicate statistically significant electrodes (Pz, P2, P4, P6, P8, PO4, PO8, O2, Oz). Colorbar represents z-values of Wilcoxon signed-rank test between high-probability cue and low-probability cue. Alpha oscillatory activity diverges in high-compared to low-probability trials over posterior electrodes in the right hemisphere. B (left) The results of a brain-to-behavioral analysis on amplitude difference between the high- and low-probability cues across a range of times (−500 to 0) and frequencies (4-40 Hz). Time 0 corresponds to stimulus onset. Black contours indicate statistically significant clusters. There is a positive correlation in a cluster centered to the alpha band: the greater the modulation of alpha activity based on prior information, the stronger the induced metacognitive bias. B (right). Topography of the significant electrodes in the brain-to-behavior analysis conducted between Δ alpha amplitude (∼7-14 Hz) extracted from each electrode (−400 to 0 ms) and metacognitive bias. The analysis revealed a cluster of electrodes located in the right parieto-occipital region (Cz, CPz, CP2, CP4, CP6, P4, P6, PO4, O2) that showed a strong association with metacognitive bias, with electrode P4 (star) exhibiting the most robust effect.

### Pre-stimulus alpha band modulation affects metacognitive bias

To assess the relationship between the behavioral modulations induced by the expectancy cue and the corresponding oscillatory dynamics, a correlation analysis was performed between participants’ metacognitive bias and the trial-type amplitude difference (high-vs. low-probability cue) at each time–frequency point within the 500-millisecond pre-stimulus window, spanning the 4–40 Hz frequency range (Figure 2B). This approach allowed directly linking individual differences in behavior with neural oscillatory activity, identifying regions in the time–frequency space where variability in oscillatory dynamics stemming from the provided prior systematically covaried with the magnitude of the metacognitive bias. The analysis revealed a significant cluster centered within the alpha band (p < 0.01), indicating that pre-stimulus alpha activity is functionally tied to the behavioral effect of probabilistic expectations observed. Specifically, a stronger prior-driven modulation of alpha amplitude corresponds to a greater metacognitive bias. To further investigate this relationship at a topographical level, a correlation analysis was conducted between Δ alpha amplitude (∼7-14 Hz) extracted from each electrode (time: –400 to 0 ms) and the metacognitive bias (Figure 2B). The analysis revealed the presence of a cluster of electrodes located in the right parietal region showing a prominent association with metacognitive bias. This confirms previous studies that showed a crucial relationship between parietal activity and metacognitive process^21,26,27^. To inform Study 2, the analysis focused to identifying the specific electrode within this cluster exhibited the strongest brain–behavior correlation, with P4 (represented by a star in Figure 2B) showing the most robust association (z value = 4.14, p < 0.001).

#### Study 2

Study 1 revealed that the expectancy cue modulates confidence judgments by inducing a metacognitive bias without altering metacognitive sensitivity. Moreover, this behavioral effect was associated with alpha activity recorded from parietal electrodes, which tracked prior-driven changes in metacognitive judgment. Therefore, a second study was conducted in which eighty-eight participants completed the same task as in Study 1, with forty-four receiving continuous Theta Burst Stimulation (cTBS) and forty-four receiving SHAM stimulation. This design allowed the investigation of the causal role of the PC in shaping metacognitive bias following prior induction (Figure 3A). It was expected that inhibiting the PC via cTBS would transiently disrupt participants’ modulation of alpha oscillations in response to probabilistic cues, impairing the integration of prior information into confidence judgments and reducing the resulting metacognitive bias.

**Figure 3.**
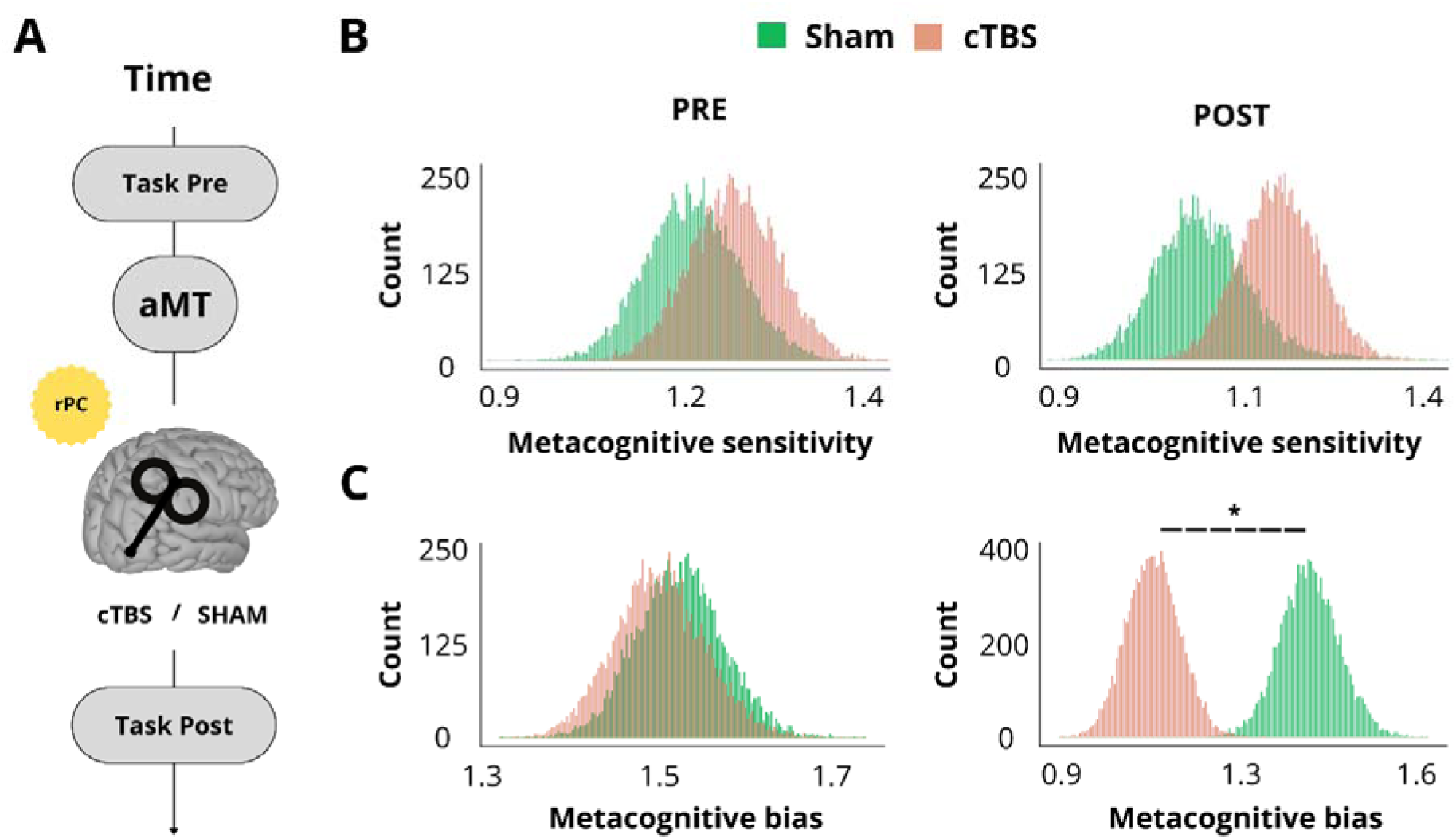
Parietal cTBS selectively reduced metacognitive bias without affecting metacognitive sensitivity. A. Experimental design. EEG data were collected during the same visual detection task used in Study 1. Participants completed three experimental blocks, after which the active motor threshold (aMT) was determined to set the stimulation intensity of the TMS protocol. Then, the continuous Theta Burst Stimulation (cTBS) or SHAM stimulation protocol was applied (N = 44 in both groups) over the right parietal cortex. After the stimulation, participants performed three experimental blocks again to test the TMS-related effect on metacognitive judgment. B. Metacognitive sensitivity (Meta-d’). Meta-d’ index is represented in PRE and POST sessions for both SHAM and cTBS conditions. The Y-axis represents the count of observations across the HMeta-d’ distribution, and the X-axis represents the most probable HMeta-d’ values. Higher Meta-d’ values indicate greater metacognitive ability. No significant changes in Meta-d’ were observed following stimulation, indicating that neither cTBS nor SHAM affected metacognitive sensitivity. C. Metacognitive bias index. Higher values indicate greater reliance on probabilistic information when making confidence judgments. The index is represented in PRE and POST sessions for both SHAM and cTBS conditions. cTBS led to a significant reduction in metacognitive bias compared to the SHAM stimulation.

### cTBS does not affect Type-I decision

The type I SDT indices *d’* and *c* were calculated to compare the effect of the expectancy cue before (PRE) and after (POST) the two types of stimulation (cTBS, SHAM; see “Supplementary Materials”). Consistent with the result of Study 1, the ANOVA revealed that the cues modulated the criterion (F_2,88_ = 81.99; p < .01; η_p_^2^ = 0.49) but not sensitivity (F_2,88_ = 1.74; p = 0.18; η_p_^2^ = 0.02). Specifically, post-hoc analysis showed that in trials where the high-probability cue was presented, participants adopted a more liberal criterion (mean ± SEM = -0.09 ± 0.06) compared to trials with the neutral-probability cue (0.29 ± 0.05; t_86_ = 8.20; p < 0.01) and the low-probability cue (0.63 ± 0.05; t_86_ = 9.56; p < 0.01), in which participants adopted a more conservative criterion relative to the neutral condition (t_86_ = 8.29; p < 0.01). Crucially, no main effects of stimulation or session were found for either *d’* or *c*, nor were there any significant interactions between stimulation, sessions, and cue (all F_2,88_ < 0.02; all p > 0.14). This result indicates that the type of cue (high-, low-, or neutral-probability) modulates the criterion but not sensitivity, independently of stimulation type (TBS, SHAM) and sessions (PRE, POST). Overall, the behavioral results indicate that inhibitory stimulation did not manipulate either the effect of the cue on response bias or perceptual sensitivity.

### Disrupting parietal cortex activity abolished cue effects on metacognitive bias

The type II SDT indices *Meta-d’*, *Meta-c and the metacognitive bias index* were calculated before (PRE) and after (POST) the two types of stimulation (cTBS, SHAM) to investigate whether the effect of the expectancy cues depends on the type of stimulation received by the subject. The analysis showed that in the cTBS group the difference in metacognitive bias between the PRE and POST sessions was significant (mean PRE = 1.53; mean POST = 1.12; mean PRE – mean POST = 0.42; HDI [0.27, 0.57]). On the contrary, in the SHAM group, the difference in metacognitive bias between PRE and POST sessions was not significant (mean PRE = 1.55; mean POST = 1.41; mean PRE – mean POST = 0.14; HDI [–0.01, 0.29]). This pointed to a reduction in the confidence bias following cTBS stimulation, due to a reduced integration of prior knowledge into metacognitive judgment (Figure 3C). Moreover, the direct comparison between the two groups showed that, in the PRE session, no differences emerged between the two groups (mean cTBS – mean SHAM = –0.02; HDI [–0.17, 0.12]). Conversely, a significant difference was observed in the POST session (mean cTBS – mean SHAM = –0.29; HDI [–0.44, –0.16]), further supporting the decrease in metacognitive bias in participants who received cTBS stimulation compared to those who received SHAM stimulation. Regarding *Meta-d’*, the analyses conducted in Study 1 showed that metacognitive performance was not affected by the cue. Therefore, to analyze metacognitive sensitivity, the cue factor was collapsed. The analysis showed the absence of significant differences in *Meta-d’* between the two groups, in both PRE (mean cTBS = 1.21; mean SHAM = 1.16; mean cTBS – mean SHAM = 0.05; HDI [–0.11, 0.19]) and POST sessions (mean cTBS = 1.15; mean SHAM = 1.07; mean cTBS – mean SHAM = 0.08; HDI [–0.15, 0.35]). Within-group comparison between the PRE and POST sessions further confirms the absence of differences (mean cTBS = 0.06; HDI [–0.10, 0.21]; mean SHAM = 0.08; HDI [–0.14, 0.36]). That implies that the type of stimulation delivered did not induce changes in the subjects’ metacognitive sensitivity (Figure3B). All in all, this evidence proved that inhibitory stimulation on the parietal area, but not SHAM stimulation, led to a reduced use of the expectation-like information when providing confidence judgments.

### cTBS affect pre-stimulus prior dependent alpha modulation

To assess the effect of cTBS on cue-based regulation of pre-stimulus oscillatory activity on posterior electrodes, we conducted a non-parametric cluster-based permutation test across the time × frequency domain in PRE and POST sessions separately for cTBS and SHAM groups (Figure 4A, 4B). Specifically, we contrasted the spectral amplitude of trials preceded by a high-probability cue against those preceded by a low-probability cue, focusing on the 500-millisecond interval before stimulus onset and covering a broad frequency range from 4 to 40 Hz. The analysis of the PRE session showed that, in both stimulation conditions, there was a suppression of alpha amplitude in trials preceded by a high-probability cue compared to those preceded by a low-probability cue (p < 0.05), mimicking the results obtained in Study 1. In the POST session however, in the cTBS group no differences in alpha amplitude were observed between the two conditions (p = 0.88), suggesting that parietal cTBS stimulation prevented participants from modulating their alpha rhythm in an expectation-driven fashion. The specificity of the stimulation is confirmed by the fact that, in the SHAM group, alpha rhythms continue to be shaped by the probabilistic cue received(p < 0.05), suggesting that participants continued to suppress alpha rhythm after the presentation of high-probability vs. low-probability cues.

**Figure 4.**
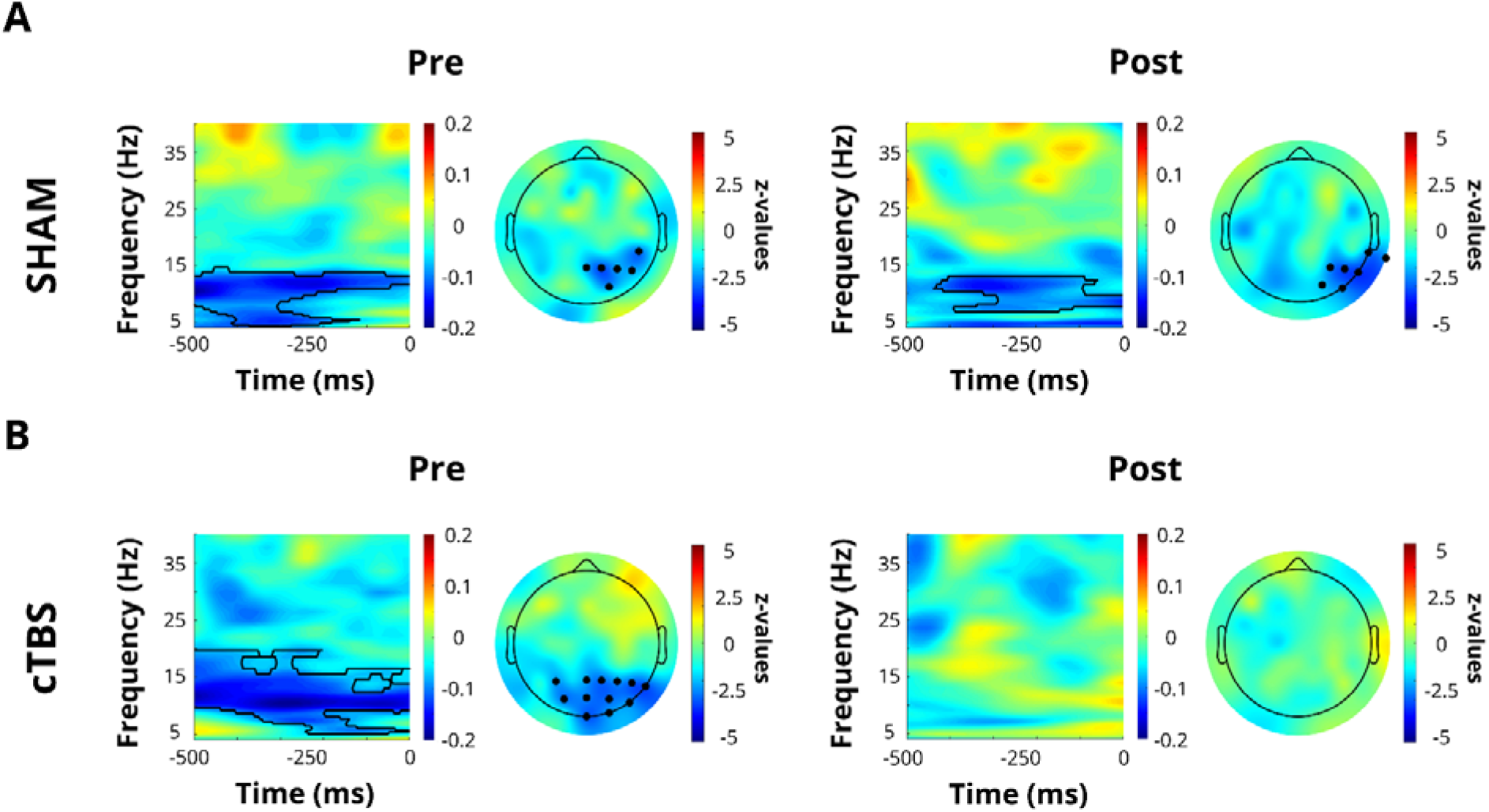
Parietal cTBS abolishes cue-induced alpha modulation. *A.* Time–frequency representations of amplitude differences between high- and low-probability cue trials at right posterior electrodes, for the SHAM group. In both PRE and POST sessions, a significant cluster centered within the alpha band was founded. Corresponding scalp topographies computed over the alpha-band cluster reveal that the effect is spatially specific to right posterior sites. In both PRE and POST sessions, a significant cluster of modulation centered within the alpha-band emerged (PRE: Pz, P4, CP6, PO4, P6, P2; POST: P4, P8, TP10, PO4, PO8, P6, TP8). *B.* Time–frequency representations of pre-stimulus amplitude differences between high- and low-probability cue trials at posterior electrodes, for the cTBS group. In the PRE session, a significant pre-stimulus cluster of modulation emerges. The topography computed confirms spatial specificity to posterior sites. (PRE: Pz, P3, Oz, O2, P4, P8, PO3, POz, PO4, PO8, P6, P2). By contrast, in the POST session no significant cluster is observed and the topography map shows no electrodes reaching significance, indicating that parietal cTBS abolished the cue-dependent modulation of the pre-stimulus alpha rhythm

Subsequently, a topographical analysis was performed to examine the spatial distribution of the observed pre-stimulus effects in PRE and POST sessions separately for cTBS and SHAM groups (Figure 4A, 4B). This involved contrasting high-versus low-probability cue trials for each electrode in an alpha frequency range (∼7-14 Hz) focusing on a 400-millisecond pre-stimulus interval. In the PRE session, the topographic cluster-based permutation test revealed a spatially specific modulation of alpha activity over parieto-occipital electrodes in both the SHAM and cTBS groups. In the POST session, the SHAM group continued to show a modulation of alpha activity over parieto-occipital electrodes, whereas no significant cluster emerged in the cTBS group. These results confirm, at a topographical level, that inhibitory parietal stimulation prevents participants from modulating their alpha rhythm in an expectation-driven fashion.

### cTBS stimulation effect on confidence is fully mediated by Alpha shift

Finally, a mediation analysis (Figure 5) was conducted to test the hypothesis that changes in alpha oscillatory activity causally mediated the relationship between stimulation conditions (cTBS vs. SHAM) and the modulation of metacognitive bias. The analysis revealed a complex pattern of relationships. The mediation pathway showed significant components: stimulation condition (cTBS vs SHAM) significantly predicted changes in alpha activity (b = –0.119, SE = 0.062, CI [–0.218, –0.027]). Most importantly, the indirect pathway from stimulation to confidence change through alpha modulation was statistically significant (indirect effect = 0.042, SE = 0.042, CI [0.001, 0.144]). Moreover, the total effect of stimulation group on confidence changes was not statistically significant (b = 0.270, SE = 0.178, CI [–0.046, 0.559]), nor was the direct effect when controlling for alpha modulations (b = 0.228, SE = 0.180 CI [–0.113, 0.504]). These findings establish alpha activity as the neurophysiological mediator through which cTBS shapes metacognitive bias.

**Figure 5.**
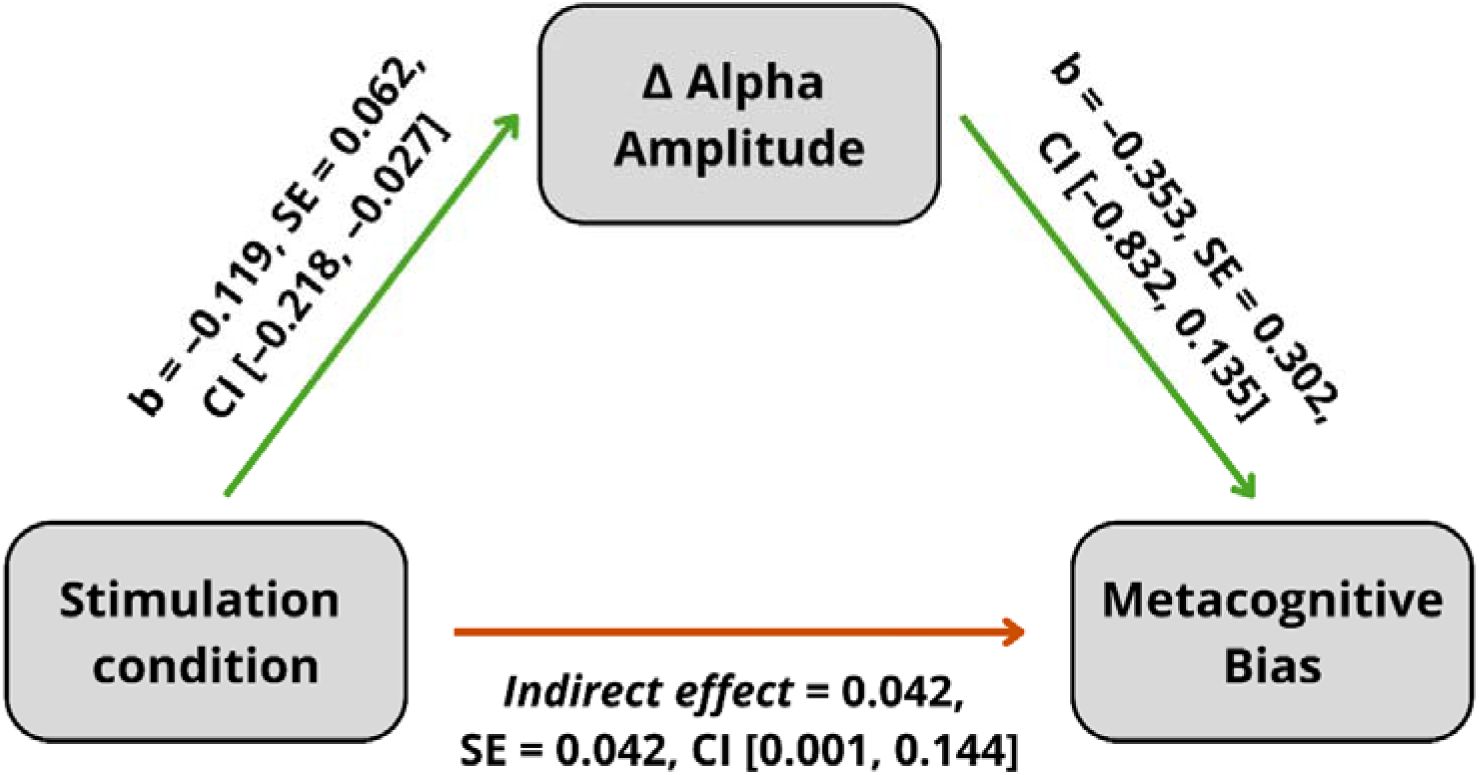
Inhibiting PC activity reduces confidence bias by concurrently modulating alpha amplitude modulation. The mediation model demonstrated that the effect of stimulation conditions (cTBS vs. SHAM) on changes in metacognitive bias is carried by modulation of pre-stimulus alpha-band amplitude. Path a show that cTBS (relative to SHAM) significantly reduces cue-dependent alpha modulation (β = –0.119, CI [–0.218, –0.027]). Crucially, the indirect effect (a_×_b) is significant (β = 0.042; CI [0.001, 0.144]), confirming that modulation of alpha amplitude fully mediates the relationship between parietal cTBS and shifts in metacognitive bias.

## DISCUSSION

Metacognition lies at the core of adaptive behavior, allowing individuals to regulate their actions by assessing confidence in their perceptual decisions. Here, in two complementary studies, we (1) detailed the electrophysiological dynamics by which prior knowledge biases metacognitive judgments and (2) established the parietal cortex’s causal contribution to this bias.

In the first study, we employed a probabilistic detection task in which symbolic cues conveyed the likelihood of target presence, combined with EEG recordings from a large sample of 75 participants. Behavioral data revealed that prior expectations significantly influenced metacognitive judgments, confirming and extending previous findings^9,11^. Specifically, prior probability cues systematically biased confidence ratings: participants expressed increased confidence when their perceptual decisions aligned with prior expectations and diminished confidence when decisions contradicted expectations. This congruency effect is consistent with a confirmation bias, whereby confidence is strongly reinforced when sensory evidence validates internal predictions. In this sense, confidence reflects a form of perceptual self-consistency: when sensory readout matches internal models, subjective certainty is reinforced; when they diverge, confidence is penalized, possibly signaling epistemic conflict between two different sources of information. Importantly, this metacognitive bias occurred without any change in metacognitive sensitivity, indicating that expectations selectively impact subjective certainty rather than metacognitive ability. Building on previous research into how prior expectations shape decision-making^7,8,15,28,29^, our findings offer a new perspective: prior knowledge does not just shape perception or choice, it also bias how people represent their own beliefs.

After investigating the metacognitive bias at the behavioral level, we characterized the neural oscillatory dynamics underpinning this effect. EEG analyses revealed a robust, expectation-driven, modulation of alpha-band amplitude: high-probability cues induced a significantly stronger pre-stimulus desynchronization than low-probability cues^8,30^. Moreover, a direct brain–behavior correlation showed that the magnitude of this alpha-band modulation predicted the degree of metacognitive bias, indicating that alpha desynchronization tracks reliance on prior beliefs during confidence judgments.

While earlier studies established that spontaneous pre-stimulus alpha amplitude predicts whether perception confidence will be high or low^13,14^, indexing fluctuations in cortical excitability^31^ that shape perceptual certainty^32^, our results uncover a qualitatively distinct mechanism. Specifically, we show that alpha activity is not only merely linked to the level of confidence, but critically to the magnitude to which confidence judgments are biased by high-level information such as probabilistic prior expectations. This transition from “confidence level” to “confidence bias” points to a more prominent functional role of alpha variability: instead of merely mirroring cortical excitability, it conditions the way internal predictions shape subjective confidence in perceptual decisions.

At the computational level, our findings fit within the metacognitive extension of SDT^33,34^. In this framework, confidence judgments depend on the magnitude of the internal response reflecting the strength of sensory evidence^35^. Confidence is assigned by comparing this response to two criteria: one for confidence in signal presence, the other for confidence in signal absence^34^. Within this model, high-confidence judgment is given when the internal response exceeds these criteria; otherwise, a low-confidence judgment is provided (Figure 6). For example, the presentation of a target inducing a strong internal response exceeds the confidence criterion and yields a high confidence judgment, while an ambiguous response would not exceed it, producing a low confidence report. Crucially, shifting the entire response distributions toward one threshold increases high-confidence judgments on that side without altering metacognitive sensitivity. Here, we propose that these shifts in response distributions originate from the cue-induced modulations of prestimulus alpha-band activity^32,36^. According to gating-by-inhibition models, alpha rhythms regulate the gain of sensory input by modulating cortical excitability: lower vs. higher alpha power increases vs. decreases excitability, thereby amplifying vs. hampering the neural representation of incoming evidence^36,37^. Consistent with this account, high-probability cues elicited a transient suppression of posterior alpha power prior to stimulus onset, indicative of increased excitability that results in a rightward shift in the internal response distributions, biasing judgments toward high-confidence “signal present” responses. In contrast, low-probability cues enhance alpha amplitude, reducing excitability and shifting the distribution toward the “signal absent” confidence boundary. Thus, fluctuations in alpha power would operate as a multiplicative gain mechanism, systematically steering the internal response distribution toward one confidence criterion or the other^32^.

**Figure 6.**
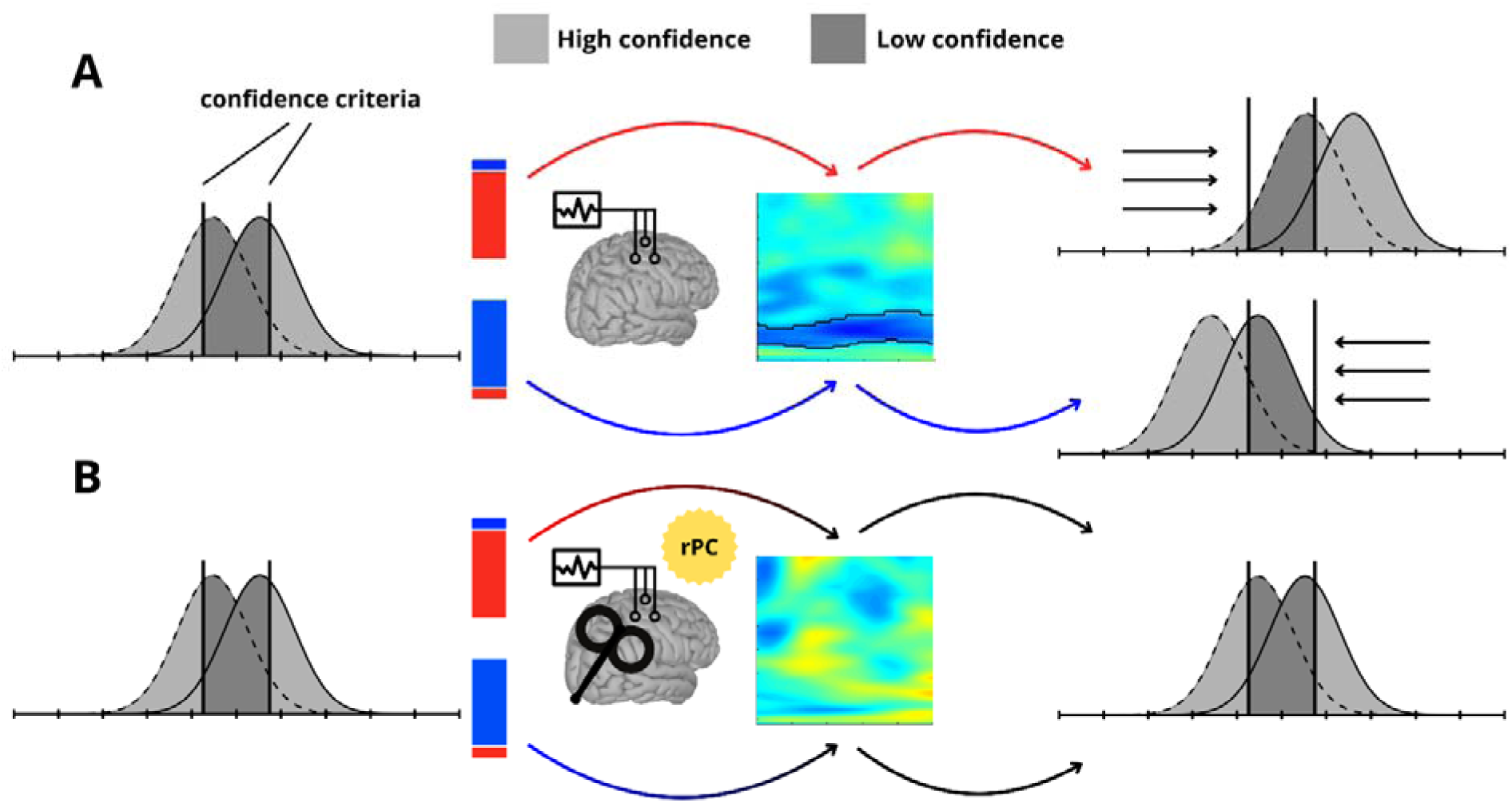
Alpha-driven shifts in the internal response distributions bias confidence. A. The figure represents the metacognitive bias induced by the cue due to alpha rhythm modulation. Within the SDT framework, a high-confidence judgment is given when the internal response exceeds the metacognitive criterion; otherwise, a low-confidence judgment is provided. As indicated by the position of the “signal absent” (dashed line) and “signal present” (solid line) distributions relative to their own metacognitive criteria, an ideal observer has an equal probability of giving high- (light grey) or low- (dark grey) confidence judgments. When the probabilistic cue is presented, there is a modulation of the alpha rhythm dependent on the information conveyed by the cue. In turn, this modulation of alpha amplitude leads to a shift in the internal distributions while the criteria remain stable. In the figure, these dynamics are illustrated by the difference in alpha amplitude between the two cues. Specifically, desynchronization (after high-probability cue) increases cortical excitability and, consequently, leads to a rightward shift in the internal response distribution. This shift results in a confirmation bias that increases the likelihood of giving a high confidence “signal present” response and decreases the likelihood of giving a high-confidence “signal absent” response. Conversely, alpha synchronization (after low-probability cue) reduces cortical excitability and, consequently, leads to a leftward shift in the internal response distribution, increasing the likelihood of giving a high confidence “signal absent” response and decreasing the likelihood of giving a high-confidence “signal present” response. B. The figure illustrates the effect on metacognitive bias following continuous theta burst stimulation (cTBS) over the right parietal cortex. After stimulation, participants showed less modulation of their alpha rhythm in response to the probabilistic cues. This lack of alpha modulation hampers the shift of internal response distributions, thereby reducing the participants’ metacognitive bias.

Interestingly, data-driven analyses showed that the relationship between expectation-driven alpha modulation and metacognitive bias was strongest over parietal channels (PC). This spatial pattern aligns with a growing body of evidence implicating the parietal cortex in the metacognitive process and with the integration of prior beliefs in perceptual decision-making^8,21,26,27,38^. For example, Park and colleagues^39^ have recently demonstrated that choice-induced confirmation biases arise not from altered sensory encoding but from a selective modulation of evidence readout in posterior parietal cortex, further pinpointing the PC as the critical hub for prior–driven biases. Our findings suggest that parietal regions contribute to confidence formation by modulating alpha activity in response to expectations. This interpretation is bolstered by previous causal evidence showing that the parietal cortex exerts control over posterior alpha rhythms^22,23^. Extending this evidence, our results offer neurophysiological support for the idea that parietal regions serve as a convergence zone where predictive and sensory signals are integrated via alpha dynamics to inform subjective certainty.

Crucially, to test the causal role of parietal alpha-mediated gain control in shaping confidence judgments, we conducted a second experiment in a cohort of 88 participants in which continuous theta-burst stimulation (cTBS) was applied over right parietal cortex (rPC)^40,41^ to transiently disrupt alpha dynamics and assess their impact on metacognitive bias. We demonstrated that inhibiting parietal activity through cTBS led to a selective attenuation of the cue-dependent modulation of pre-stimulus alpha oscillations. Strikingly, this neurophysiological disruption was mirrored in behavior: participants’ metacognitive bias induced by expectation cues was attenuated. Specifically, the characteristic congruency effect observed in Study 1, where confidence increased when perceptual responses aligned with expectations and decreased when they did not, was reduced.

Our findings provide causal evidence that parietal activity is not merely correlated with, but necessary for, embedding prior beliefs into metacognitive evaluation. In addition, mediation analysis confirmed that disruptions in alpha dynamics due to cTBS mechanistically mediated the observed changes in metacognitive judgments. This effect was specific to active stimulation and absent in the SHAM group, confirming its causal specificity. Within the SDT metacognitive framework introduced above, these results further suggest that, by disrupting the alpha-mediated gain mechanism, cTBS abolished the shifts in the internal response that give rise to metacognitive bias. As a result, confidence ratings become less sensitive to prior information, reducing the metacognitive bias. This result could be also framed within the predictive coding framework^1^, with this effect reflecting a breakdown in precision-weighted inference: following parietal activity suppression, alpha dynamics become less sensitive to probabilistic cues weakening the influence of expectations and shifting confidence toward bottom-up evidence.

The selective reduction in metacognitive bias following cTBS, without concomitant changes in metacognitive sensitivity, mirrored the pattern observed in our first study and aligned with theoretical expectations. This finding is also in line with recent causal evidence showing that inhibitory neurostimulation of the parietal cortex selectively disrupts spatial choice bias during endogenous attention tasks^42^. Although their focus was on attentional reorienting, the pattern of bias modulation closely parallels our metacognitive results. Taken together, these studies converge on the notion that the PC serves as a domain-general hub for bias regulation, spanning both spatial orienting and confidence judgments. However, it is conceivable that, under different conditions, where prior knowledge affects both confidence and metacognitive sensitivity^9^, the parietal cortex would contribute to integrating expectations into both components. In this framework, the parietal cortex would operate as a dynamic weighting system, calibrating the relative impact of expectations on metacognitive processes according to the task at hand.

Together, these findings reveal for the first time that metacognitive judgments are not passively read out from sensory evidence but are actively constructed through the integration of internal predictions. We show that prior beliefs shape confidence via modulations of posterior alpha rhythms, and that disrupting parietal activity with cTBS causally abolishes this bias. This selective, oscillation-based mechanism revealed how the brain dynamically gates expectation-driven confidence. More broadly, our findings suggest that the regulation of neural excitability via oscillatory dynamics gives rise to a metacognitive confirmation-bias aimed at preserving perceptual self-consistency. Specifically, parietal alpha activity mediated this process: expectations, once consistently matched with sensory readout, tend to solidify into beliefs, as we seek coherence between what we anticipate and what unfolds in our perception. Within this framework, we propose that alpha oscillations contribute to the maintenance of cognitive homeostasis by adaptively tuning sensory excitability to align with contextual information. This regulatory role underlines the brain’s ability to maintain perceptual consistency in uncertain environments, offering a mechanistic entry point to understand, and ultimately modulate, confidence computations in both healthy cognition and neuropsychiatric conditions.

## METHODS

### Participants

#### Study 1

Seventy-five healthy participants (39 female) took part in the study. This sample is drawn from a previously published dataset^15^.

#### Study 2

Eighty-eight healthy participants (56 female) took part in the study. Participants were randomly assigned to the experimental group and the control group and received the cTBS (44 participants) or SHAM (44 participants) protocol, respectively. Both groups were matched for numerosity and gender.

In both studies, participants were aged between 18 and 35 years. All participants provided informed consent before taking part in the study, which was conducted in accordance with the Helsinki Declaration and received approval from the Bioethics Committee of the University of Bologna (protocol code 201723, approved on 26 August 2021). None of the participants had a clinical history of neurological or psychiatric disorders, nor any conditions that would contraindicate the use of TMS (according to Rossi and colleagues^43^).

### Stimuli and Experimental task

The stimuli were presented on an 18’ CRT monitor (Cathode Ray Tube, CRT) with a display resolution of 1280 × 1024 pixels and a refresh rate of 85 Hz. They consisted of checkerboards presented in the lower left visual field (Figure 1A). Each cell of the checkerboards could contain small gray circles (target) or not (catch trials). The stimuli were created and presented using MATLAB (version 2016, The MathWorks Inc., Natick, MA) in combination with the Psychophysics Toolbox^44^.Participants were instructed to indicate, as quickly and accurately as possible, the presence or absence of gray circles within the checkerboard using the “K” and “M” keys on the keyboard. Additionally, they were asked to respond with their right hand. Throughout the task, participants were instructed to maintain their gaze on a fixation cross displayed at the center of the screen.

Each participant completed an adaptive titration procedure to determine the contrast of the gray circles (perceptual threshold) required to achieve a target detection accuracy of ∼70%. The overall accuracy in the main task was approximately 70% (namely, 71.9%), confirming the effectiveness of the titration phase. This procedure was conducted by presenting to the participant an equal number of trials with the target present and trials with the target absent (catch trials). The latter were included to prevent potential confounding effects caused by response bias; in fact, presenting only trials with the target present would not allow us to determine whether differences in threshold values were due to actual perceptual ability or to a different decision-making style^45^.

During the test phase (FIGURE 1A), each block consisted of 90 trials. At the beginning of each trial, a cue indicating the probability of target presence was displayed at the center of the screen for 1 second, followed by a fixation cross. The cue consisted of a rectangle with its lower portion colored red and the upper portion colored blue. The percentage of red indicated the probability of the target being present. Cue high and cue low (informative cues) indicated the probability of the presence of the target of 67% and 33%, respectively. Instead, the neutral cue (un-informative cue) equally predicted (50%) the presence and absence of the target. The probability of target presentation was in accordance with the one indicated by the cue. Participants were also explicitly told that the probabilistic cue was congruent with the actual probability of stimulus presentation. After a variable interval of 1.2–1.5 s, a checkerboard appeared in the lower-left quadrant of the screen (with or without the gray circles) for 60 ms. At this stage, the contrast of the gray circles was set to the threshold intensity identified through the adaptive titration procedure. The stimulus was presented in the right hemifield to prevent interference in the results due to spontaneous variations in attention between hemifields. After the participant provided their response, they were asked to indicate their confidence level regarding the response just given. Confidence was rated using the keyboard keys “Q,” “W,” “E,” and “R,” each corresponding to a different confidence level (Q: not confident at all, W: slightly confident, E: moderately confident, R: highly confident).

### Experimental procedure

Participants were seated in a dimly lit room in front of a monitor positioned at a distance of ∼ 57 cm. The EEG cap was then fitted according to the international 10-10 system. Initially, participants were familiarized with the task through a training phase, and each participant’s individual perceptual threshold was determined using an adaptive titration procedure.

#### Study 1

Participants were required to complete 6 blocks of the experimental task (Figure 1A).

#### Study 2

Participants completed 3 experimental blocks, after which the active motor threshold^46^ was determined and the cTBS or SHAM stimulation protocol was applied, with participants being randomly assigned to one of the protocols. Finally, 5 minutes after the stimulation^47^, participants performed 3 experimental blocks again (Figure 3A).

### Type-I Signal-detection theory (SDT) modeling

The SDT measures *d’* and *c*^6^ were computed. *d’* quantifies a participant’s stimulus sensitivity, with higher values indicating greater discriminative ability. In contrast, *c* quantifies a subject’s decision criterion. A *c* value different from 0 indicates the presence of a response bias. Specifically, a *c* value greater than 0 reflects a more conservative criterion (i.e., tendency to give “absent response”), while a *c* value smaller than 0 reflects a more liberal criterion (i.e., tendency to give “present response”).

### Type-II Signal-detection theory (SDT) modeling

The Type-II measures *Meta-d*’ and *Meta-c*^33,34^ were computed. *Meta-d’* is a measure of metacognitive sensitivity expressed within the framework of Type-I SDT. It can be understood as the amount of signal available to the observer for performing the confidence evaluation task (Type-II task). The higher the *Meta-d’*, the greater the participant’s metacognitive ability. An individual with optimal metacognitive sensitivity will always be more confident when correct and less confident when incorrect.

Confidence criteria (*Meta-c*), on the other hand, represent Type-II bias calculated within the *Meta-d’* framework and indicate the tendency to give high or low confidence ratings. A *Meta-c* value close to 0 suggests a more liberal criterion (i.e., tendency to give high confidence ratings), while a *Meta-c* value farther from 0 indicates a more conservative criterion (i.e., tendency to give low confidence ratings).

### Continuous Theta Burst Stimulation

Participants received either continuous Theta Burst Stimulation (cTBS) or SHAM stimulation. The type of stimulation each participant received was randomly assigned. The stimulation was delivered using a Magstim rapid TMS machine with an eight-shaped coil (diameter = 7 cm) and consisted of triplets of pulses at a 50 Hz frequency, administered every 200 ms (5 Hz) for a total duration of 40 seconds, resulting in 600 pulses overall. This procedure induces inhibitory effects on cortical excitability lasting ∼50 minutes after stimulation^40^. For each subject, the entire POST session was conducted within this time window. The stimulation intensity was individualized for each participant and set at 80% of their active motor threshold (aMT). The aMT was defined as the minimum stimulation intensity (expressed as a percentage of the maximum output power of the Magstim device) that elicited a visible muscle contraction in the right hand in at least 3 out of 5 trials when a single pulse was applied to the left motor area. During this procedure, the participants maintained a slight muscle contraction in the hand opposite to the stimulation side. For technical and safety reasons, the maximum stimulation intensity was set at 45% of the device’s maximum output power. The mean stimulation intensity was 43.5% of the maximum output power. The stimulation was applied over electrode site P4 in line with the results from the brain-to-behavior analysis conducted in Study 1, which showed that P4 is the electrode showing a higher correlation between Metacognitive bias modulations and alpha amplitude modulations (see results of Study 1 for details). For SHAM stimulation, the same protocol was followed, but the coil was positioned perpendicularly to the scalp to prevent any actual neurophysiological effects. Participants were asked to indicate the stimulation condition they thought they had received (active, SHAM, or uncertain); a chi-square test revealed no significant differences between groups (X^2^ = 0.83, p = 0.66), indicating that participants could not reliably distinguish between active and SHAM conditions.

### EEG preprocessing and time-frequency decomposition

#### Study 1, Study 2

Participants comfortably sat in a room with dimmed lights. A set of 64 electrodes was mounted according to the international 10–10 system. EEG signals were acquired at a rate of 1000 Hz and all impedances were kept below 10 kΩ. EEG was processed offline with custom MATLAB scripts (version R2022b) and with the EEGLAB toolbox^48^. The EEG recording was filtered offline in the 0,5–100 Hz band and a notch-filter at 50 Hz was applied. The signals were visually inspected, and noisy channels were spherically interpolated. Epochs spanning −4100 to 2000 ms relative to checkerboard onset were extracted and individual trials were visually checked and those containing excessive noise, muscle or ocular artefacts discarded. Next the recording was then re-referenced to the average of all electrodes, and the Independent Component Analysis (ICA) was applied, an effective method largely employed for removal of EEG artefacts. Components containing artifacts that could be clearly distinguished from brain-driven EEG signals were subtracted from the data. After these steps, the signals were downsampled to 256 Hz and a Laplacian transform was applied to the data using spherical splines. The Laplacian is a spatial filter that aids in topographic localization by attenuating artifacts attributable to volume conduction, rendering the data more suitable for performing connectivity analyses^49^. Subsequently, time-frequency analysis was implemented by convolving the time series data with a set of complex Morlet wavelets (whose cycles increased between 5 and 13 cycles as a function of frequency), defined as complex sine waves tapered by a Gaussian. Convolution was performed via frequency-domain multiplication, in which the Fourier-derived spectrum of the EEG data was multiplied by the spectrum of the wavelet, and the inverse Fourier transform was taken. Then, the phase was obtained by extracting the angle relative to the positive real axis and the amplitude by extracting the absolute value of the resulting complex time series. Amplitude was then condition-specific baseline-corrected using a decibel (dB) transform: dB amplitude = 10 × log10 (amplitude/baseline). Baseline amplitude was defined as the average amplitude in the period ranging from −3100 to −2700 ms before stimulus onset.

### SDT Type-I analysis

#### Study 1

To examine the influence of the probabilistic cue on sensitivity and criterion, *d’* (Z(Hit rate)−Z(False Alarm rate)) and *c* (−1/2 * [Z(Hit rate)+Z(False Alarm rate)]) were calculated separately for trials preceded by high-, low-, or neutral-probability cues (Figure 1B). Repeated measures ANOVA were conducted with cue type as a within-subject factor (three levels: high-, low-, and neutral-probability) to statistically assess cue-related effects on sensitivity and criterion. Post-hoc paired-sample t-tests were performed to interpret the ANOVA results, with p-values corrected for multiple comparisons.

#### Study 2

To investigate potential differences related to stimulation received *d*’ and *c* were computed separately for trials preceded by high-, low-, or neutral-probability cues, considering both stimulation protocol (cTBS, SHAM) and sessions (PRE, POST; see “Supplementary Materials”). To examine these effects, a three-way repeated-measures ANOVA was conducted with cue type (three levels: high-, low-, neutral-probability) and sessions (two levels: PRE, POST) as within-subject factors, and stimulation condition (two three levels: cTBS, SHAM) as a between-subject factor. Post-hoc paired-sample t-tests were performed to interpret the ANOVA results, with p-values corrected for multiple comparisons.

### SDT Type-II analysis

#### Study 1

To evaluate the effect of the probabilistic cue on metacognitive sensitivity and confidence criteria, *Meta-d’* and *Meta-c* were calculated separately for trials preceded by a cue indicating high-, low-, or neutral-probability of target presence (Figure 1C, 1E). The *Meta-d’* and *Meta-c* values were extracted using the Hierarchical *Meta-d’* approach (*HMeta-d’*)^25^. The analysis was performed using the hierarchical fitting option available in the *HMeta-d’* toolbox. This approach involves a Bayesian hierarchical estimation of the parameters, which returns the most probable value of the parameter in each condition. Specifically, this method integrates Bayesian priors to constrain estimates of both the group’s average metacognitive sensitivity and individual efficiency using hierarchical modeling. The Bayesian hierarchical estimation implemented in *HMeta-d’* specifies prior density distributions at the group level for each participant-level parameter and provides a group-level estimate of metacognitive sensitivity (*Meta-d’*). Group-level fits were performed using the function *fit_meta_d_mcmc_group*.

To analyze the effect of the cue on metacognitive sensitivity, *Meta-d’* was examined by first computing the distribution of differences in the posterior parameter samples for each cue (high-, low- or neutral-probability cue) and then determining the 95% Highest Density Interval (HDI) for this distribution. The group-level posterior densities were then used to test the statistical significance of differences in metacognitive sensitivity. Specifically, if the 95% HDI of the difference between conditions did not include 0, the difference was considered credible, whereas if the HDI included 0, the difference was deemed non-significant.

Subsequently, to analyze the effect of the probabilistic cue on *Meta-c*, the same methodology was used as in the previous analyses. *Meta-c* was calculated for high-, low-, or neutral-probability cues for type-I responses (present/absent) separately, resulting in three distinct *Meta-c* values. Finally, to examine the effect of the cue on *Meta-c* a general index called Metacognitive bias was computed (Figure 1D). This index was derived by summing the difference between the *Meta-c* value in absent responses in trials preceded by low-probability cue (LpA) and in trials preceded by high-probability cue (HpA), with the difference between the *Meta-c* value in present responses in trials preceded by low-probability cue (LpP) and in trials preceded by high-probability cue (HpP).

(*Metacognitive bias = [Meta - c_LpA_ - Meta-c_HpA_] + [Meta-c_LpP_ - Meta-c_HpP_])*. A greater value suggests a larger shift in bias for congruent responses vs. incongruent ones.

#### Study 2

To investigate whether the effect of the probabilistic cue on metacognitive sensitivity and confidence criteria differed depending on the stimulation condition, *Meta-d’* and *Meta-c* were calculated separately before and after the two stimulation conditions (cTBS, SHAM; Figure 3B, 3C). The analyses conducted in Study 1 showed that cognitive performance was not affected by the cue. Therefore, to analyze metacognitive sensitivity, the cue factor was collapsed, and the distributions of the cTBS and SHAM conditions, obtained through the hierarchical model, were compared in the PRE and POST stimulation sessions. An overlap of the distributions within the 95% HDI was considered indicative of a non-significant difference.

Subsequently, to analyze the presence of differences in *Meta-c*, the Metacognitive bias was calculated as described in Study 1, both in the PRE and POST sessions for both groups (cTBS, SHAM). Next, the difference in the distributions of the Metacognitive bias between the PRE and POST sessions was calculated separately for each group. Values above 0 indicate a greater Metacognitive bias in the PRE compared to the POST session, while negative values indicate a greater Metacognitive bias in the POST compared to the PRE session. Additionally, to analyze the difference of the cue’s effect on the Metacognitive bias between the two groups before and after stimulation, the difference between the cTBS and SHAM conditions was calculated in the PRE (cTBS_pre_ - SHAM_pre_) and in the POST (cTBS_post_ - SHAM_post_). Negative values indicate a difference between the two groups in the Metacognitive bias. Finally, the difference between the PRE and POST sessions was calculated in order to directly compare the two groups (i.e. [cTBS_pre_ - SHAM_pre_] and [cTBS_post_ - SHAM_post_]). If the 95% HDI of the difference did not include 0, the difference was considered statistically significant, whereas if the interval includes 0, the difference was deemed non-significant. Additionally, a value greater than 0 indicates a greater decrease in the use of the prior in the POST compared to the PRE session, specifically in the cTBS group.

### EEG analysis - Oscillatory amplitude

#### Study 1

To assess whether there is relationship between pre-stimulus oscillatory activity and the behavioral modulations induced by the expectancy cue, an amplitude analysis on a cluster of central and right posterior electrodes was conducted, given that the visual stimuli were presented in the left visual field (Figure 2A). Specifically, the mean amplitude was computed across the following electrodes: O2, PO4, PO8, P2, P4, P6, P8. This approach helped minimize potential biases related to the selection of a limited number of sensors. To examine pre-stimulus oscillatory activity associated with prior utilization, a non-parametric permutation test based on a time × frequency cluster analysis (n = 1000) was performed on the amplitude difference between high- and low-probability trials, within a 500-millisecond pre-stimulus window, considering a broad frequency range (4-40 Hz). For each permutation, trial types were shuffled within each participant, generating a null distribution of amplitude differences. This data-driven method allowed us to test, point by point, for significant differences between the two prior conditions across the entire time window and for all included frequencies while controlling for multiple comparisons^50^. Subsequently, a topographical analysis was performed to examine the spatial distribution of the observed pre-stimulus effects. This involved contrasting high-versus low-probability cue trials for each electrode in the alpha frequency range (∼7-14 Hz) within a 400-millisecond pre-stimulus interval. A topographical cluster-based permutation was performed (Figure 2A), applying the non parametric Wilcoxon signed rank test^51^ to analyze spatially specific modulation of alpha activity over right parieto-occipital electrodes. For each permutation trial types were shuffled within each participant with the same shuffle for each electrode.

#### Study 2

To assess whether the relationship between pre-stimulus oscillatory activity and the behavioral modulations induced by the expectancy cue differed based on the stimulation condition, the same EEG analysis of Study 1 was conducted separately for the two stimulation conditions (cTBS, SHAM) in both sessions (PRE, POST; Figure 4A, 4B).

### EEG analysis - Brain-to-behavior

#### Study 1

To investigate the relationship between the behavioral modulations induced by the expectancy cue and the oscillatory modulations, a correlation analysis was conducted between metacognitive bias and the trial-type amplitude difference (high- vs. low-probability) at each time–frequency point within a 500-millisecond pre-stimulus window, across the 4-40 Hz band using MATLAB’s *robustfit* function (Figure 2B). This analysis allows outliers to be preserved while assigning them less weight, ensuring more reliable estimates in the presence of non-ideal data. Each point in the time-frequency map obtained from the previously described analysis was used as a predictor and correlated with the metacognitive bias. Time–frequency points with correlation values not surviving the statistical threshold (p > 0.05) were removed, and clusters were subsequently defined as groups of adjacent time–frequency points exhibiting significant correlations. Then a correction for multiple comparisons was run using a cluster-based permutation approach. For each permutation the metacognitive bias was shuffled between each participant generating a null distribution of behavioral patterns. Finally, a topographical analysis was performed by extracting, for each electrode (time: –400 to 0 ms), the Δ alpha amplitude (∼7-14 Hz; i.e., the amplitude difference between high- and low-probability trials in the alpha band) and correlating it with the metacognitive bias (Figure 2B). The alpha shift was selected as the variable of interest as the cluster emerged from the previous data-driven analysis that predicted behavioral performance was centered in the alpha range. This approach allowed us to identify the spatial distribution of the correlation across the scalp and to determine the electrode where the relationship was strongest. To investigate this, we performed robust regressions between the prestimulus oscillatory amplitude difference (high vs. low probability) and individual metacognitive bias at each electrode. For each regression, we extracted the t-statistic of the slope (i.e., the estimate divided by its robust standard error). The electrode showing the largest absolute t-statistic was considered to exhibit the strongest correlation. Finally, a correction for multiple comparisons was run using a cluster-based permutation approach.

#### Study 2

To further test the role of alpha oscillatory activity in driving the effect of stimulation type on changes in metacognitive bias, we conducted a one tail mediation analysis using PROCESS for JASP^52^, specifying Model 4 (Figure 5). The analysis was tested one-tailed, based on the evidence that, following cTBS, there was a significant reduction in cue-related alpha modulation. Since alpha activity was positively correlated with metacognitive bias, it was expected that its suppression would be accompanied by a corresponding decrease in bias. The experimental group was coded as a binary predictor (0 = SHAM, 1 = TBS), while both the mediator (Δ alpha amplitude) and the dependent variable were computed as pre-to-post stimulation differences (Δ Metacognitive bias). Indirect effects were estimated using bias-corrected bootstrap resampling with 1000 iterations. The analysis yielded estimates of the direct effect (c’), indirect effect (a × b), and total effect (c) of the stimulation group on confidence shift. Significance of the mediation pathway was assessed based on whether the bootstrap confidence interval for the indirect effect excluded zero in the hypothesized direction.

## Data Availability

Data available to corresponding authors upon reasonable request.

## Competing interests

The authors declare no competing interests.

## Acknowledgements

VR is supported by Next Generation EU (NGEU) and funded by the Ministry of the University and Research (MUR), National Recovery and Research Plan (NRRP) PRIN 2022 (grant n 2022H4ZRSN - CUP J53D23008040006): Predictive waves in human perception and individual differences along the autism-schizophrenia continuum (D DN. 104 02.02.2022); (grant n P2022XAKXL – CUP J53D23017340001): Investigating the plasticity of human predictive coding through neuromodulation (D DN. 1409 14.09.2022); Ministerio de Ciencia, Innovación y Universidades, Spain (PID2019-111335 GA-100); Bial Foundation (033/22)

## Contributions

Conceptualization: L.T., V.R.

Methodology: L.T., D.R., A.P.

Investigation: L.T., M.C., D.R., A.P.

Visualization: L.T., A.P., D.R.

Supervision: VR., L.T.

Writing—original draft: L.T., A.P., D.R., V.R.

Writing—review & editing: V.R., M.C., L.T, A.P., D.R.

## Notes

### Competing Interest Statement

The authors have declared no competing interest.

